# Ileal mucus viscoelastic properties differ in Crohn’s disease

**DOI:** 10.1101/2024.01.03.574052

**Authors:** Catharina Kramer, Hanna Rulff, Jörn Felix Ziegler, Nadra Alzain, Annalisa Addante, Aditi Kuppe, Sara Timm, Petra Schrade, Philip Bischoff, Rainer Glauben, Julia Dürr, Matthias Ochs, Marcus A. Mall, Michael Gradzielski, Britta Siegmund

**Author notes:** Correspondence:* Britta Siegmund, MD, Department of Gastroenterology, Infectiology and Rheumatology, Charité - Universitätsmedizin Berlin, Hindenburgdamm 30, 12200 Berlin, Germany, |. These authors have contributed equally to this work and share first authorship (Catharina Kramer, Hanna Rulff).

## Abstract

Crohn’s disease (CD) is an inflammatory bowel disease (IBD) that can affect any part of the gastrointestinal tract, frequently involving the terminal ileum. While colonic mucus alterations in CD patients have been described, terminal ileal mucus and its mechanobiological properties have been neglected. Our study is the first of its kind to decipher the viscoelastic and network properties of ileal mucus.

With that aim, oscillatory rheological shear measurements based on an airway mucus protocol that was thoroughly validated for ileal mucus were performed. Our pilot study analyzed terminal ileum mucus from controls (n=11) and CD patients (n=11). Mucus network structure was visualized by scanning electron microscopy (SEM).

Interestingly, a statistically significant increase in viscoelasticity as well as a decrease in mesh size was observed in ileal mucus from CD patients compared to controls. Furthermore, rheological data were analyzed in relation to study participants clinical characteristics, such as inflammatory status, revealing noteworthy trends.

In conclusion, this study provides the first data on the viscoelastic properties and structure of human ileal mucus in the healthy state and in CD, demonstrating significant alterations between groups and highlighting the need for further research on mucus and its effect on the underlying epithelial barrier.

## 1. Introduction

Crohn’s disease (CD) is a disease entity of inflammatory bowel disease (IBD). It is a chronic transmural disease that can affect any part of the gastrointestinal tract, but the terminal ileum is frequently involved. The disease is characterized by chronic inflammation, and in subpopulations fistulizing and stricturing disease can occur, resulting in a profound overall reduction in quality of life ^1^. Although a number of factors in the pathogenesis of the disease have been identified, a causal treatment is still lacking ^1^. Within the pathogenesis, the intestinal barrier plays a key role and shows significant differences between the ileum and the colon ^2^. Whilst the majority of data in the mucus literature has focused on the colon, the focus of this study will be on the ileum. The ileum is formed by the epithelial cell layer, which represents epithelial cells in different states of differentiation as well as Paneth cells that produce antimicrobial peptides (AMPs). The ileum is covered by a single layer of mucus composed of mucins, AMPs and Immunoglobulin A and is permeable to nutrient uptake and microbiota ^3, 4^. While there is considerable data on the contribution of epithelial dysfunction to disease pathogenesis (reviewed in detail in ^5^), the function of the mucus layer in human ileal CD in particular requires further attention.

The mucin (MUC) composition of ileal mucus has been described previously and includes the membrane-tethered mucins MUC3, MUC13, MUC17 ^6^ and the secreted mucins MUC2, MUC5AC, MUC5b and MUC6 ^6, 7^. Data on mucus in the ileum of CD patients are limited and suggest a downregulation of MUC3, MUC4 and MUC5B during inflammation ^8^. In addition, mucus contains AMPs, lysozyme, and various lectins ^9, 10^. In a subgroup of CD patients, Paneth cell dysfunction has been identified as a central mode of disease, leading to reduced antimicrobial activity and increased abundance of adherent invasive *Escherichia coli* (AIEC) ^11^. However, more detailed analyses of human ileal mucus are currently lacking. CD is a chronic inflammatory disease that leads to structural changes in the intestinal wall, with the development of strictures, resulting in changes in tissue mechanics. Recent data indicate that changes in tissue mechanics may also represent a novel form of danger signal ^12^ and directly affects the interaction with immune cells ^13^. Therefore, a better understanding of the role of mucus and the biophysical and biochemical triggers contained in the mucus layer during ileal inflammation compared to the healthy state is needed.

Mucus can be described as a complex hydrogel consisting of more than 95% water and other components such as glycoproteins, lipids, proteins, DNA, and salts ^14^. A complex interaction of mucus components is responsible for its proper functioning. Mucin glycoproteins play an essential role in this ^15, 16^. Mucins are high molecular weight macromolecules with extensive glycosylation and a high water-binding capacity. Therefore, in a solubilized form, they have the properties of a complex fluid ^17–19^. Mucins can form a three-dimensional multimeric network with a pronounced mesh-like structure. This mesh-like structure is responsible for the versatile biophysical properties of mucus, such as its crucial filtering function ^20–23^. Mucus can adapt to different physiological conditions, but its proper functioning can be severely altered and cause severe multi-organ disease as exemplified by the rare genetic disease cystic fibrosis (CF) ^22, 24, 25^.

Mucus is a viscoelastic gel with the capacity to both flow (viscous element) and deform (elastic element). The viscoelastic behavior is determined by the network architecture and the complex interplay among its constituents, which enable mucus to perform its essential functions ^16, 26, 27^. The viscoelastic properties of intestinal mucus are critical in preserving its vital attributes, such as protective functions, efficient lubrication, barrier formation, and adaptability to varying conditions.

The terminal ileum is of particular interest in the pathogenesis of CD. This is not only due to the significant increase in microbiota density compared to the upper small intestine, but also because it serves as a critical site for the priming of the immune system ^2^. Mucosal homeostasis depends on a functionally intact intestinal barrier. The luminal border of this barrier is formed by the mucus layer, which serves as a critical defense against pathogens while facilitating the absorption of various nutrients ^28^. Although the composition, especially the presence of antimicrobial peptides, has been studied previously, there is a lack of information regarding the biophysical properties of the small intestinal mucus.

To the best of our knowledge, this study is the first to analyze the viscoelastic properties of human intestinal mucus and to compare the healthy mucus to that of CD patients.

The viscoelastic properties of mucus in the terminal ileum can be determined through systematic macrorheological measurements. In this study, dynamic oscillatory shear measurements such as an amplitude sweep, and strain-controlled frequency sweep were conducted using a cone-plate geometry. By obtaining the viscoelastic parameters, it is possible to determine the extent of modifications of the mucus properties in a diseased compared to the healthy state. Additionally, we used scanning electron microscopy (SEM) images to improve understanding of the mucus network structure and any possible differences between mucus from controls and CD patients. By combining these methods, our objective was to gain a more exhaustive comprehension of the internal, functional characteristics of mucus in the terminal ileum.

## 2. Methods

### 2.1 Study design and participants

This study was approved by the Ethics Committee of the Charité - Universitätsmedizin Berlin (EA4/120/20). All participants gave written informed consent to participate in this study. Patients between 18 and 60 years of age were included. Patients underwent colonoscopy at the Department of Gastroenterology, Infectiology and Rheumatology, Charité - Universitätsmedizin Berlin. The indication for colonoscopy was to determine disease activity in Crohn’s disease (CD) patients and to screen for colorectal cancer in the control group. Patients at high risk for colonoscopy due to anticoagulation, severe cardiovascular or pulmonary disease, and patients with a history of ileal surgery were excluded.

The demographic and clinical characteristics of the study participants [controls (n=11) and CD patients (n=11)] are shown in **Table 1**. As the study was designed as a pilot study, patients irrespective of CD activity were included.

**Table 1:** Demographic and clinical characteristics of study participants. Demographic and clonical characteristics of control and Crohn’s disease patients. Abbreviations: TNF-α = tumor necrosis factor alpha, IL = interleukin, n.d. = not determined, HBI = Harvey-Bradshaw Index, y = years; GI = gastrointestinal

|  | Controls | Crohn's disease |
| --- | --- | --- |
| Age, mean years (min-max) | 45.58 (20-56) | 33.73 (20-46) |
| Sex (male/female) | 3/8 | 4/7 |
| Montreal classification | n.d. |  |
| A1 (<17y) |  | 2 |
| A2 (17-40y) |  | 8 |
| A3 (>40y) |  | 1 |
| L1 ileal |  | 0 |
| L2 ileocolonic |  | 1 |
| L3 colonic |  | 10 |
| L4 upper GI tract |  | 3 |
| B1 non-stricturing non-penetrating |  | 6 |
| B2 stricturing |  | 1 |
| B3 penetrating |  | 4 |
| p perianal disease |  | 3 |
| Disease duration, mean years (min-max) |  | 12.5 (1-30) |
| Smoking |  |  |
| yes | 3 | 4 |
| no | 8 | 7 |
| Medication |  |  |
| Anti-TNF-α | | 7 |
| Anti-IL-12/IL-23 |  | 2 |
| Antimetabolites | 1 | 1 |
| none |  | 2 |
| Clinical inflammation mean (HBI) | n.d. | 4.55 (0-19) |
| Endoscopic inflammation |  |  |
| none | 11 | 8 |
| mild | 0 | 2 |
| severe | 0 | 1 |
| Histologic inflammation |  |  |
| 0 | 2 | 5 |
| 1-3 | 2 | 2 |
| >4 | 0 | 2 |
| n.d. | 7 | 2 |

In all patients, disease activity was determined on clinical, endoscopic, and histological level. Clinical disease activity was defined as a Harvey Bradshaw Index (HBI) greater than 5. To determine inflammation at the sample site, specifically the terminal ileum, the Simple Endoscopic Score (SES-CD), a well-established metric commonly applied in colonoscopy, was used. A SES-CD score of 0-2 in the terminal ileum was considered uninflamed, while scores of 3 6 indicated mild inflammation and a score of 7 or higher indicated severe inflammation ^29–31^. The modified Naini and Cortina score was used to assess histologic inflammation by an experienced pathologist who was double-blinded ^32^. Furthermore, all patients completed the same questionnaire to provide information on their disease course and dietary habits.

### 2.2 Sample collection and preparation

Patients prepared for colonoscopy by taking laxatives the day before, according to standard protocols. During colonoscopy, native mucus was carefully collected from the terminal ileum by using a lavage catheter (ENDO-FLEX GmbH, Voerde, Germany). Mucus samples were aliquoted immediately after collection. Aliquots used for rheological measurements were stored at 4 °C. Samples were measured either the same day after resting for two hours following colonoscopy or the next day.

The following exclusion criteria were applied to ensure sample quality and the reliability of the rheological measurements: For sample quality, samples that appeared macroscopically diluted or visibly contained stool or blood were excluded. For rheological measurements, samples with a non-linear amplitude sweep or low G’ and G’’ values in the same range as the laxative solution measurements were excluded. As a consequence, 10 (5 controls and 5 CD patients) of the original 32 samples had to be excluded. The remaining n=11 controls and n=11 CD patients were included in the analysis of their macrorheological parameters.

### 2.4 Macrorheological measurements

We used the previously described macrorheological protocol for human airway mucus ^33^. Details are provided in the *Supplementary Information*. In brief, the oscillatory rheological shear measurements were performed at 37 °C using a solvent trap to ensure saturation of the atmosphere at all times during the experiment. The amplitude sweep was performed at a frequency of 1 Hertz (Hz) and a shear deformation γ between 0.01-10%. The following frequency sweep was conducted at a shear deformation of 2% and covered a frequency range of 0.5-50 Hz. The macrorheological data were analyzed focusing on the viscoelastic moduli, the storage modulus G’ and the loss modulus G’’.

### 2.5 Sample preparation for scanning electron microscopy

For scanning electron microscopy (SEM), native mucus from the terminal ileum of 3 CD patients and 3 controls was immersed in 2.5% glutaraldehyde (Serva, Heidelberg, Germany) in 0.1 M cacodylate buffer (Serva, Heidelberg, Germany) immediately after sample collection. After this first fixation, the samples were postfixed with 1% OsO_4_ (Electron Microscopy Sciences, Hatfield, USA) in 0.1 M cacodylate buffer for 2 hours at room temperature (RT) and stored overnight at 4 °C in 0.1 M cacodylate buffer. To ensure a good solution exchange between all steps while minimizing sample loss, samples were resuspended in each solution and then centrifuged (500 g). The supernatant was always removed without drying the samples. 30 µl of the pellet was applied to a poly-d-lysine coated coverslip. After sedimentation, the samples were dehydrated in a graded ethanol series. To completely dry the samples while preserving the surface structure as much as possible, the samples were critical point dried (Leica EM CPD300, Leica, Wetzlar, Germany). After mounting the coverslips on SEM pin stubs, sputtering with gold palladium, and loading into the SEM (GeminiSEM300, Zeiss, Oberkochen, Germany), the samples were examined by meandering scanning of the coverslips with 7 kV high voltage and a secondary electron detector.

### 2.7 Statistical Analysis

Data analysis was performed with GraphPad Prism version 10.1.1 (GraphPad Software, San Diego, CA, United States). The results were reported as mean ± standard error of the mean (SEM). A Kruskal-Wallis test followed by Dunn’s multiple comparison and a Mann-Whitney test were used to test for statistical significance. Statistical significance was defined by p < 0.05.

## 3. Results

### 3.1 Demographic and clinical characteristics of study participants

The study population consisted of 11 controls and 11 Crohn’s disease (CD) patients. **Table 1** shows the detailed demographic and clinical characteristics of the study participants. There was a higher proportion of female subjects. Since this was a pilot study, the selection of CD patients was primarily based on the presence of CD, rather than a specific subtype within the Montreal Classification. The degree of disease activity was assessed applying the Harvey-Bradshaw Index. Additionally, endoscopic inflammation was classified based on the SES-CD score, and histologic inflammation was categorized using the modified Naini and Cortina score, as detailed in the *Methods* section.

### 3.2 Validation of rheological measurements of intestinal mucus

During colonoscopy, mucus was gently collected from the epithelial cell layer of the terminal ileum as depicted in **Figure 1A**. The mucus was evaluated according to the established criteria explained in the *Methods* section and rheologically measured. Colonoscopy is not possible without prior cleansing of participants with laxative solution. To distinguish between native mucus and laxative solution in this study, we validated our measurements by comparing the macrorheological data. Laxative solutions did not exhibit viscoelastic properties resembling those of a gel, as demonstrated in **Figure S1**. In contrast, rheological behavior of the native ileal mucus clearly showed a hydrogel character and stable measurements for each ileal mucus sample from controls and CD patients (**Figure S2**).

**Figure 1:**
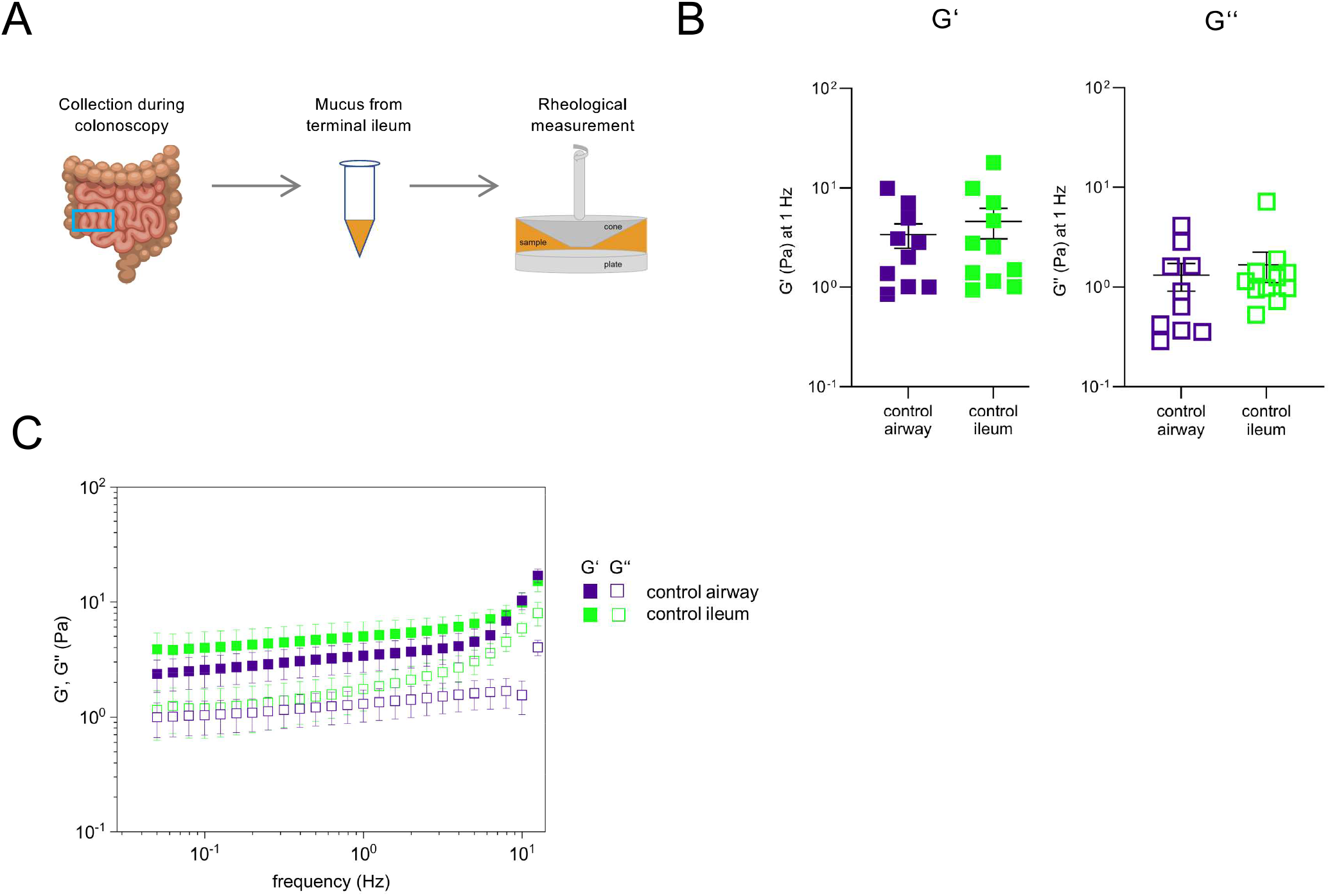
Rheological characterization of ileal mucus compared to airway mucus. (**A**) Illustration of the experimental workflow: obtaining sample from the terminal ileum via endoscopy, sample aliquotation, rheometer. (**B**) Comparison of airway and ileal mucus from controls.. Mean storage modulus G’ and loss modulus G’ at a frequency of 1 HZ. (**C**) Storage modulus G’ and loss modulus G’’ of airway mucus and ileal muscus from controls as function of frequency. N=10 airway mucus from controls (blue squares) and n=11 control ileal mucus (n=11) (green squares). Data are shown as mean ± standard error of the mean (SEM). All measurements were done at 37 °C.

As the samples were measured either on the same or the following day after storage at 4 °C overnight, we compared those groups. We found no difference in the viscoelastic properties between ileal mucus samples measured on the same day of colonoscopy and samples measured one day later, indicating that this is not a cause for concern (**Figure S3**).

To our knowledge rheological data on human intestinal mucus are absent; therefore, we compared our findings on human ileal mucus from controls (n=11) with macrorheological data on human airway mucus from controls (n=10; **Figure 1B**). Some of the airway mucus samples from controls were previously included in a publication ^33^. The demographic and basic clinical characteristics of the control group for airway mucus are presented in **Table S1**. The macrorheological protocol used to determine the viscoelastic properties of airway mucus from controls provided robust and reliable results ^33^. It has been widely used to analyze the viscoelastic properties of human airway mucus from controls and changes in patients with cystic fibrosis. Airway mucus macrorheology proves to be a highly promising biomarker for monitoring disease severity and progression, and for assessing treatment response ^34, 35^. The airway mucus from controls exhibited a viscoelastic solid behavior, typical of mucus hydrogel ^21, 35–38^ with a storage modulus G’ five times larger than the loss modulus G’’ (**Figure 1B**). Interestingly, the viscoelastic moduli G’ and G’’ of ileal mucus from controls were found to be in a similar range to those of the airway mucus from controls. The mean storage modulus G’ at 1 Hz in airway mucus from controls was 3.4 Pascal (Pa) compared to 4.7 Pa in ileal mucus from controls. Additionally, as in airway mucus from controls, the storage modulus G’ (mean = 4.6 Pa) was greater than the loss modulus G’’ (mean = 1.7 Pa) in ileal mucus from controls indicating hydrogel behavior.

**Figure 1C** shows the mean values (+/− SEM) of the viscoelastic moduli at 1 Hz of the airway mucus and ileal mucus from controls over the entire frequency studied. The stability of both viscoelastic moduli could be observed for both groups. The detailed values for G’ and G’’ of the n=10 airway mucus samples from controls over the investigated frequency range are shown in **Figure S4**.

### 3.3 Higher viscoelastic moduli in ileal mucus of Crohn’s disease patients

In **Figure 2A**, the viscoelastic moduli of the ileal mucus from controls and CD patients are presented as mean values, indicating a significant difference between the two groups. This difference is evident throughout the entire frequency range, with the storage module G’ exceeding the loss modulus G’’ in both groups. At the representative frequency of 1 Hz, both the storage modulus G’ as well as the loss modulus G’’ of ileal mucus from CD patients showed higher values than those of ileal mucus from controls (**Figure 2B**). The difference in viscoelastic moduli between ileal mucus from controls and CD patients is about a factor of 15-20 (absolute differences: 95 Pa for G’ and 40 Pa for G’’). Upon detailed analysis of the frequency sweeps of the ileal mucus from controls and CD patients (**Figure 2A/C/D**), it is evident that the mucus from CD patients exhibited a relatively more frequency-independent behavior. Specifically, the storage modulus G’ and the loss modulus G’’ remained consistent across the frequency range investigated. It is worth noting that the ratio of G’/G’’ slightly increased with increasing frequency in the ileal mucus from the CD patients but decreased in the ileal mucus from the controls (**Figure S2**). This suggests that ileal mucus from CD patients is relatively more elastic at higher frequencies compared to controls. This also shows that stable measurements with reliable values for the viscoelastic moduli of both groups were performed (**Figure 2A**, **Figure S2**). Furthermore, it can be observed that there is a greater heterogeneity in the viscoelastic parameters of the ileal mucus from CD patients within this group when compared to the controls (**Figure 2C/D**).

**Figure 2:**
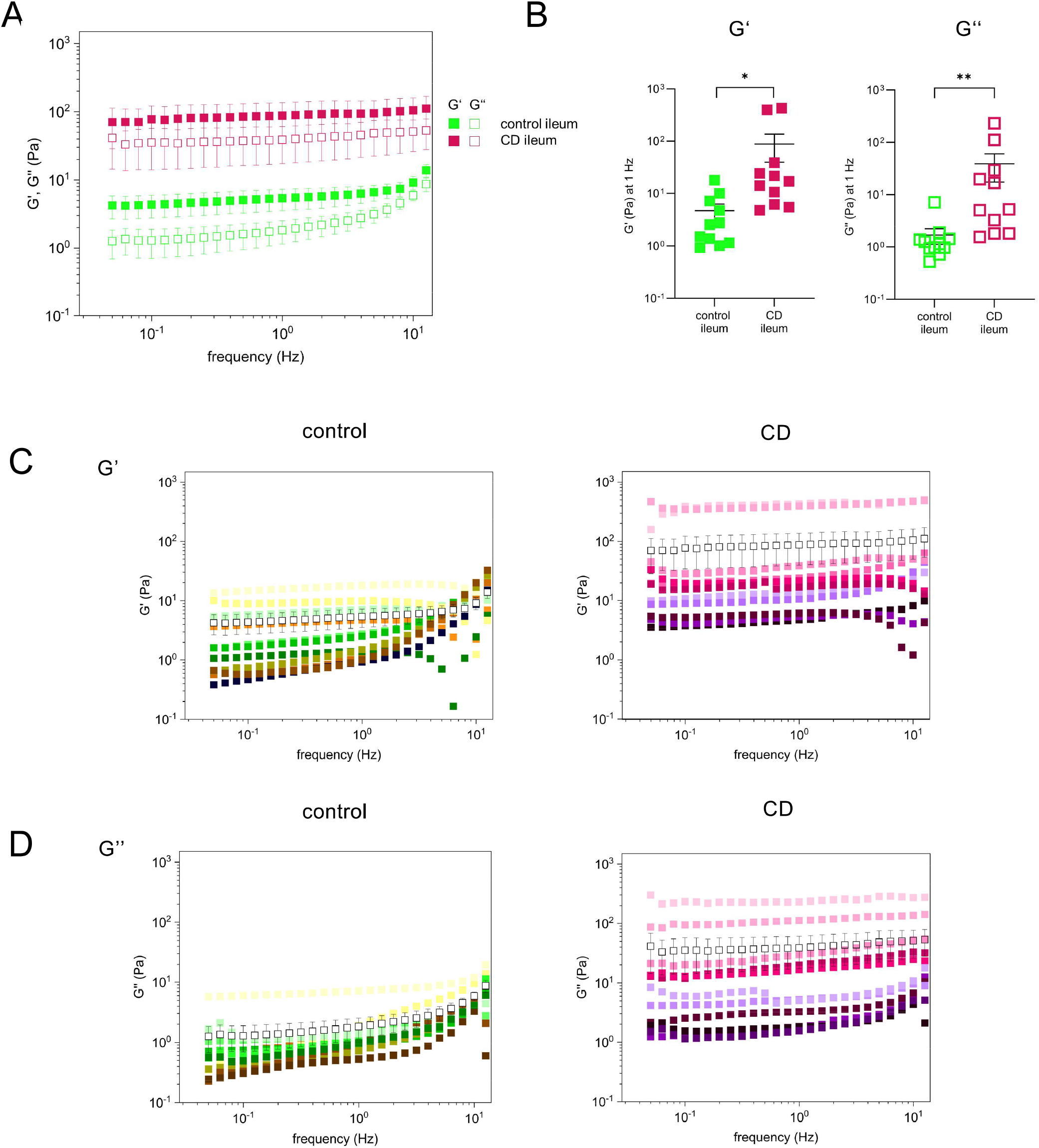
Viscoelastic properties of ileal mucus from Crohn’s disease (CD) patients compared to controls. (**A**) Mean storage modulus G’ and loss modulus G’’ of ileal mucus from controls and from CD patients as function of frequency. Data are shown as mean ± standard error of the mean (SEM). (**B**) Storage modulus G’ and loss modulus G’’ of ileal mucus from controls and CD patients at a frequency of 1 Hz. Data are shown as individual values and mean ± standard error of the mean (SEM). (**C**) Storage modulus G’ and (**D**) loss modulus G’’ of ileal mucus from controls and CD patients as function of frequency. Data are shown as individual values (colored) and mean ± standard error of the mean (SEM) (open black squares with error bars); *p < 0.05, **p < 0.01, ***p < 0.005; always n=11 ileal mucus from controls (green squares) and n=11 ileal mucus from CD patients (red squares).

### 3.4 Rheological data related to different clinical characteristics

In the next step, we evaluated whether there was a correlation between the clinical characteristics of the patients and the viscoelastic moduli G’ and G’’ (**Figure 3**). The exact mean values can be taken from **Table S2** of the *Supplementary Information*.

**Figure 3:**
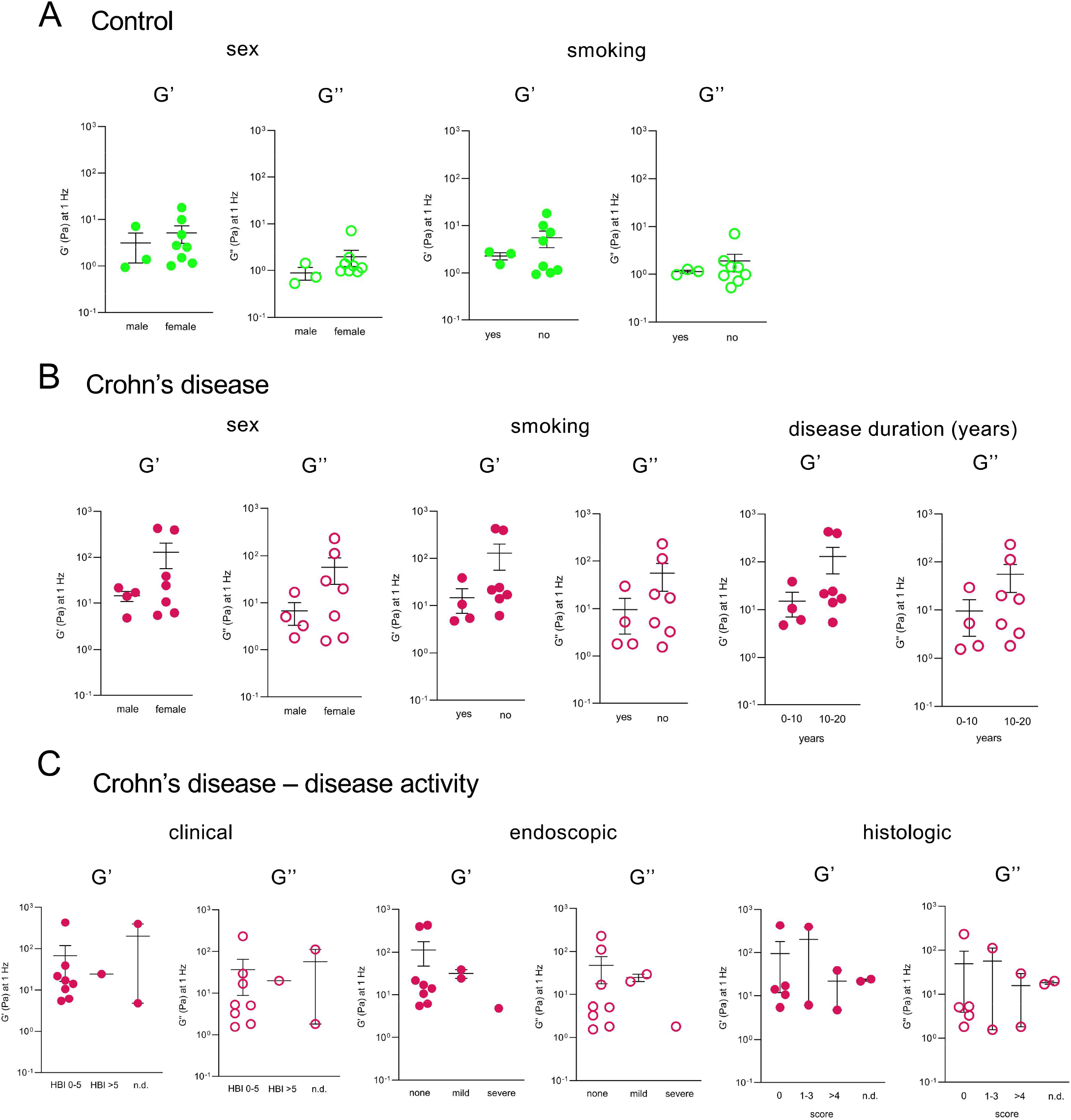
Rheological data in relation to clinical characteristics. Storage modulus G’ and loss modulus G’’ of ileal mucus were measured at a frequency of 1 Hz. (**A**) Shown is ileal mucus from controls (n=11; green circles) and (**B**) Crohn’s disease CD) patients (n=11; red circles) divided depending on sex (male/female), smoking (yes/no), and duration of CD (years). (**C**) Ileal mucus from CD depending on clinical (according to HBI), endoscopic (according to SES-CD; 0-2 = none, 3-6 = mild, >7 = severe inflammation) and histologic inflammation (according to modified Naini and Cortina score) status at the time of colonoscopy is presented. Data are shown as individual values and mean ± standard error of the mean (SEM). Abbreviation: n.d. = not determined.

When looking at sex and smoking behavior, the majority of controls as well as CD patients were predominantly female and non-smokers. Interestingly, females and non-smokers tended to have higher mean values for G’ and G’’ compared to males and smokers, consistently across groups (**Figure 3A/B**).

Within the group of CD patients, it is interesting to note that those with a disease duration of 10-20 years had a higher mean value for G’ and G’’ than those with a disease duration of 0-10 years (**Figure 3B**). The observed tendency cannot be attributed to differences in age among the patients, as there were no significant variations in G’ and G’’ across different age groups. This observation also holds true for the control group (**Figure S5**).

In the following, CD patients were grouped according to clinical, endoscopic, and histologic disease activity as classified by HBI, SES-CD, and modified Naini and Cortina scores as described in the *Methods* section (**Figure 3C**). Noticeably, the patients with low endoscopic inflammation also showed low histological inflammation. In this pilot study, there were no significant differences in the mean values of G’ and G’’ between patients with different levels of inflammation, possibly due to the small group size and the small number of CD patients with higher levels of inflammation.

### 3.5 Structural characterization by scanning electron microscopy indicates a tighter network for Crohn’s disease ileal mucus

The rheological data (**Figure 2B**) allow to estimate an effective mesh size ξ (nm) (**Figure 4A**), which is inversely proportional to the shear modulus and can be envisioned to be related to the mean width of the pores in the mucus network. The estimated mesh size of the ileal mucus from CD patients (mean value = 60 nm) was significantly smaller than that of the ileal mucus from controls (mean value = 120 nm). This indicates a tighter meshwork with smaller pore size and less permeability in the case of CD. Notably, similar mesh size differences were observed in airway mucus; the mesh size of airway mucus from patients with cystic fibrosis is approximately 82 nm and that from controls approximately 125 nm, resulting in a difference of 43 nm ^33, 39^.

**Figure 4:**
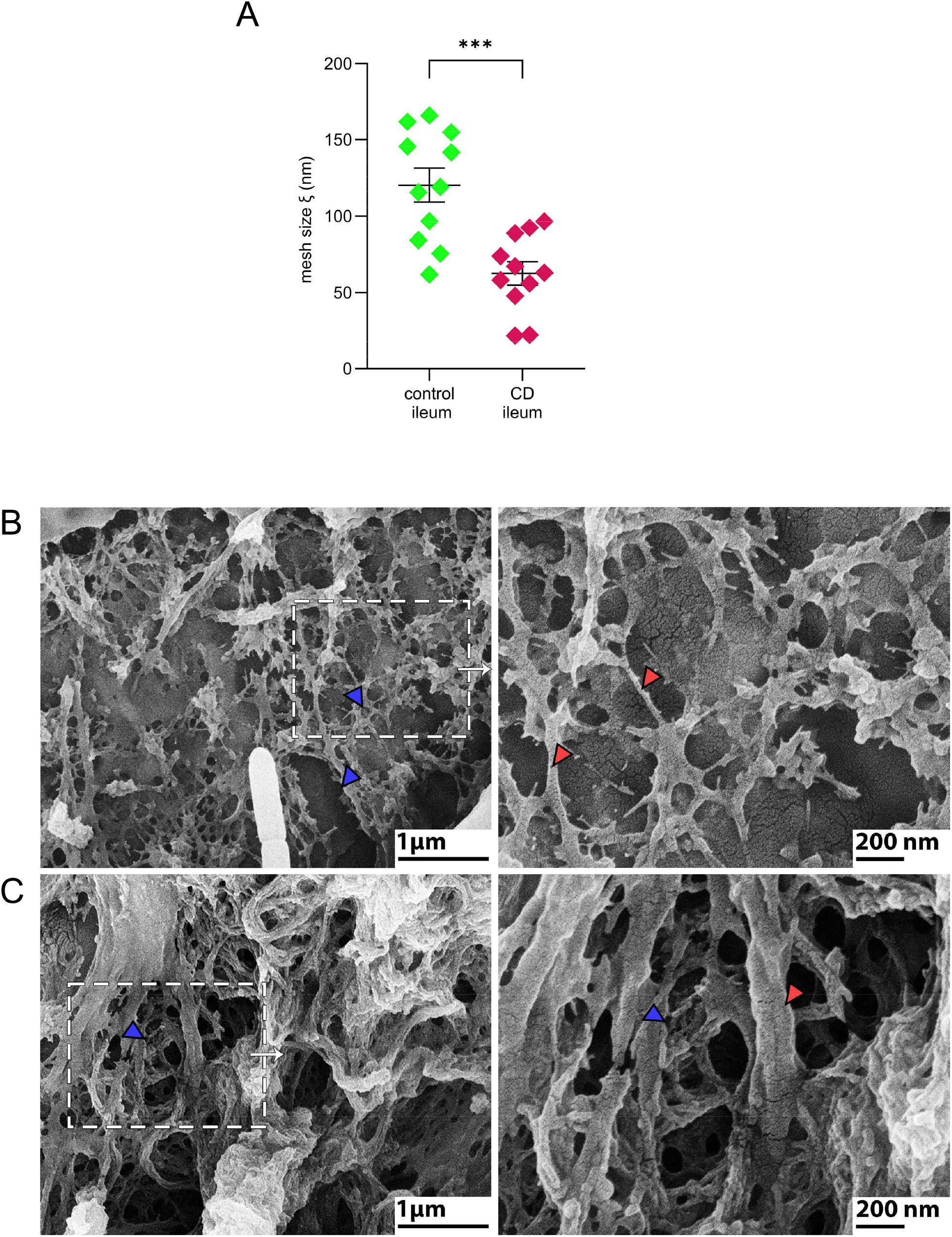
Estimated mesh size for ileal mucus in controls and Crohn’s disease (CD) patients and scanning electron microscopy. (**A**) Effective mesh soze ξ (nm) of ileal mucus from controls (n=11; green squares) and from CD patients (n=11; red squares) calculated from rheological data. Data are shown as mean ± standard error of the mean (SEM); *p < 0.05, **p < 0.01, ***p < 0.005. **Images in two magnification scales are shown with boxed areas on the left being depicted enlarged on the right.** (**B**) Representative area of mucus from a control revealing a loose structure, pores of different sizes (blue arrowheads) and thin filaments (red arrowheads). (**C**) Representative area of mucus from a Crohn’s disease (CD) patient. The network shows a compact, interwoven organization with many smaller pores (blue arrowheads) and partially very thicker filaments (red arrowhead).

However, rheology allows only a rough estimate of such mesh sizes and therefore we employed SEM to obtain more detailed structural information on the different mucus samples. Representative images are shown in **Figure 4B, 4C**. In both, the control and CD patient ileal mucus, the mucus exhibited a filamentous organization with numerous pores. The control samples revealed a loosely structured mesh characterized by thin filaments and varying pore sizes. In some cases, large pores could be observed. In comparison, mucus from CD patients displayed a more compact, interwoven filament arrangement, characterized by thicker filaments and many smaller pores.

## 4. Discussion

This study is the first to systematically investigate viscoelastic and network properties of human small intestinal mucus of controls and patients with crohn’s disease (CD) using macrorheology and scanning electron microscopy. First, we were able to show that the well-established rheological protocol for airway mucus can equally be applied to intestinal mucus. Of note, the viscoelastic properties of mucus from the terminal ileum of the control group demonstrated similarities to the properties of airway mucus from control patients. Interestingly, the ileal mucus from CD patients showed higher viscoelastic moduli and a tighter network structure compared to that of controls.

As the analysis of the viscoelastic properties of human intestinal mucus using macrorheology is a novelty, we first had to establish a protocol. For this purpose, we utilized a protocol that had been validated for airway mucus from both controls and patients with cystic fibrosis (CF) (**Figure 1, Figure S4**) ^33^. Applying this protocol to our terminal ileum mucus samples, we observed robust and reliable rheological data (**Figure 1**, **Figure S2**). Samples that were diluted with laxative solutions could be safely excluded due to their substantially altered rheological properties (**Figure S1**). Furthermore, our studies show that ileal mucus samples can be stored at 4 °C for 24 h before measurement without changes in the viscoelastic properties (**Figure S3**).

Remarkably, our data are the first to reveal differences in the viscoelastic moduli and the network properties of the ileal mucus between controls and CD patients. While the viscoelastic moduli G’ and G’’ for ileal mucus of controls were comparable to airway mucus, the values for G’ and G’’ were approximately one decade higher in CD patients compared to controls. For the ileal mucus of controls as well as CD patients G’ dominated over G’’ over the entire frequency range. This means that the elastic part of the system is larger than the viscous part for both groups (**Figure 2**). From the rheological data, the mesh size of the mucus network can be calculated. The pore diameter of ileal mucus was approximately 60 nm smaller in CD patients, indicating a more densely packed mucus network compared to controls (**Figure 4A**).

To confirm our findings and further characterize the mucus network with its detailed structure with an additional method, we examined our samples using SEM (**Figure 4B/C**). SEM is well suited to visualize internal structures on a microscopic level and has been successfully used for the investigation of mucus samples on several occasions ^40, 41^. The compact and interwoven filament network with smaller pores observed in ileal mucus of CD patients confirmed the rheological findings in comparison to controls. Thus, the functional differences are reflected by ultrastructural changes.

The lack of available data on the viscoelastic properties of human terminal ileum and colon mucus poses a significant challenge for interpreting our results. To address this issue, we turned to human airway mucus, which is well-studied, particularly in the context of inflammatory lung disease such as CF ^42^. By comparing the two, we developed hypotheses about the mechanisms that may contribute to the observed differences in viscoelastic properties between CD patients and controls.

It is known that in airway diseases like CF, inflammation plays a crucial role in altering the biophysical properties of mucus. The hypothesis that inflammation can induce crosslinking within the mucus’s internal structure, such as the formation of disulfide bonds and other crosslinking types, provides a compelling explanation for the higher viscoelastic moduli of airway mucus from patients with CF compared to controls^25, 43^. Crosslinking in mucus, similar to that observed in cystic fibrosis, may also play a significant role in altering the properties of mucus in CD.

The dehydration of airway mucus in inflammatory airway diseases like CF is linked to altered ion and fluid transport mechanisms which can lead to mucus with altered viscoelastic properties impacting its proper functioning ^44^. It can be hypothesized that the inflammatory mechanism in CD could induce the ion and fluid transport in the terminal ileum as a compensatory mechanism for the excessed mucus crosslinking explaining the altered viscoelastic properties in ileal mucus from CD patients showing elevated inflammation scores (**Figure 3C**). In line with this hypothesis are previous data indicating that electrogenic sodium absorption via ENaC is impaired in CD colon, even in the absence of inflammation ^45^. Furthermore, epithelial tight junctions determine paracellular water and ion movement. Claudin-2 forms a channel for small cations and water and is typically expressed in small intestine. It builds a major pathway for the paracellular transport of sodium, potassium, and fluid. Remarkably, claudin-2 is up-regulated in CD ^46, 47^.

The capacity for fluid storage in CD mucus is reduced compared to healthy mucus, which contrasts with increased fluid release during inflammation ^48^. The reduced fluid storage capacity may result in a less effective barrier ^49, 50^, leading to increased susceptibility to infection and autoinflammation. During inflammation, the observed increase in fluid release may be a compensatory mechanism that attempts to restore mucus clearance and pathogen removal. The observed changes in viscoelastic properties between ileal mucus from CD patients and ileal mucus from controls could be caused by the different composition of mucus in healthy and diseased state.

The abnormal ileal mucus properties in CD could lead to significant pathophysiological consequences. They may disrupt the normal transport of ileal content, affecting nutrient absorption and bowel motility. Additionally, changes in mucus could alter interactions with the gut microbiota, potentially disturbing the microbial balance and triggering inflammation ^3, 51^. This inflammation could further exacerbate organ dysfunction, creating a feedback loop of worsening symptoms and systemic effects.

After general comparison of rheological data of controls and CD patients the analysis was extended in a further step by taking into account the clinical characteristics of the study participants (**Figure 3**, **Table 1**). Interestingly, some trends could be identified. It is noticeable that the patients with a higher disease duration (10-20 years) had approximately one-decade higher values for G’ and G’’ than the patients with a lower disease duration (0-10 years). This suggests that duration of disease affects the composition of the intestinal epithelial cells and hence mucus composition. Also, when looking at sex, age, and smoking, we could observe some differences that were consistent in controls as well as CD patients. Due to the small cohort size in this pilot study, those differences were not statistically significant and need to be addressed in further studies.

Limitations of our study are due to the small cohort sizes. As our aim was to test the method in a pilot study, certain imbalances in the cohorts can be noted. For example, there are more women than men in our study. Also, in the control group the participants were older on average, since their reason for colonoscopy was cancer surveillance. Furthermore, it can be noted that not many of the CD patients in our study did show a high inflammation status (**Table 1**). When there is a lot of inflammation, the terminal ileum becomes highly inflamed and swollen. That leads to difficulties collecting sufficient amounts of mucus through the catheter in the terminal ileum or to exclusion of the samples due to blood contamination.

Nevertheless, our present cohort is well characterized; its clinical and demographic parameters provide the basis for further analyses and follow-up studies. For example, a detailed analysis of defined patient subgroups such as non-stricturing versus stricturing ileal disease as well as focusing on gender differences or smoking behavior.

In conclusion, this study provides the first data on the viscoelastic properties of human ileal mucus or even human intestinal mucus in general. Furthermore, our data indicate significant changes in CD mucus compared to mucus of controls in terms of viscoelastic properties, but also in terms of calculated mucus mesh size and mesh structure. This initial analysis warrants for detailed follow-up analyses to correlate mucus composition and viscoelastic properties and the subsequent effects on the underlying epithelial barrier.

Additionally, the rheological method that was used here is also applicable to monitor and evaluate the evolution of airway diseases and the success of therapeutic options in CF patients ^39^, hence can potentially offer a similar novel monitoring tool in CD.

## 6. Data Availability

All data are provided in the original manuscript. The raw data will be made available by the authors, without undue reservation.

## 7. Ethics statement

This study was approved by the Ethics Committee of the Charité - Universitätsmedizin Berlin (EA4/120/20).

## 8. Author contribution

Conception and design of the study: HR, CK, JFZ, RG, NA, MG, BS, MAM

Acquisition, analysis, and interpretation of data: HR, CK, JFZ, MG, BS; MAM, MO, ST, JD, AA, AK, PS, PB

Drafting the article or revising it critically for important intellectual content: HR, CK, JFZ, RG, MG, MO, JD, BS, MM, AA, AK, ST, PS, PB

All authors contributed to the article and approved the submitted version.

## 9. Acknowledgements

The authors thank the patients with crohn’s disease and the controls for their participation in this study. The authors thank the Core Facility for Electron Microscopy of the Charité for support in acquisition of the data. This study was funded by the Deutsche Forschungsgemeinschaft (DFG, German Research Foundation; Project ID 431232613) – SFB 1449 projects A01, B01, B04, C04, Z02 and the Germany Federal Ministry of Education and Research (82DZL009B1 to M.A.M.). P.B. and J.F.Z. are participants in the BIH Charité Clinician Scientist Program funded by the Charité -Universitätsmedizin Berlin, and Berlin Institute of Health (BIH). P.B.’s work is additionally funded by a DKTK Berlin Young Investor Grant 2022.

## Supplementary information

### Methods

#### Demographic and basic clinical characteristics of controls providing airway mucus

**Supplementary Table 1:** Demographic and basic clinical characteristics of controls providing airway mucus. Abbreviations: BMI = body mass index; FEV1 = forced expiratory flow in one second; SEM = standard error of the mean; n.d. = not determined.

| | Controls<br>Mean (± SEM)<br>or n (%) |
| --- | --- |
| Number of samples | 10 |
| Age, years | 33.6 (± 1.6) |
| Sex, female | 80 % |
| BMI, kg/m<sup>2</sup> | 26.0 (± 1.4) |
| FEV<sub>1</sub> % predicted | n.d. |

### Macrorheological measurements

A Kinexus Pro+ Rheometer (NETZSCH GmbH, Selb, Germany) with a stainless-steel cone-plate geometry with a cone-diameter of 20 mm and a cone-angle of 1° was used. The sample was transferred to the lower geometry with a non-electrostatic spatula. After confining the sample with the upper rotating cone, the spatula was also used to trim any excess sample. A passive solvent trap was used and a defined amount of 900 µL distilled water was added into the upper reservoir of the solvent trap. The water filled solvent trap ensured that the atmosphere in this closed environment was saturated at any time during the experiment while measuring at 37 °C. After loading and between the performed sequences the samples were equilibrated for 5 minutes at the given temperature to ensure full temperature equilibration and sufficient network relaxation. Each sequency included an amplitude sweep and a frequency sweep down- and upwards. The amplitude sweep was performed at a fixed frequency of 1 Hz and covered a range of shear deformation γ between 0.01-10%. The frequency sweep was conducted at a fixed shear deformation of 2% and covered a frequency range of 0.5-50 Hz. An entire measurement sequence lasted around 40 minutes.

The linear viscoelastic (LVE) region was determined within the amplitude sweep. The LVE represents the quasi-static behavior of the material commonly at lower strains. The frequency sweeps were analyzed regarding the behavior of the storage modulus G’ and loss modulus G’’.

To estimate the effective mesh size ξ of the mucin network, we used the following formula:

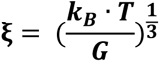

with k_B_ being the Boltzmann constant, T the absolute temperature and G the shear modulus, which we approximated by the G’ value at 1 Hz.

### Results

#### Validation of rheological measurements of intestinal mucus

**Supplementary Figure 1:**
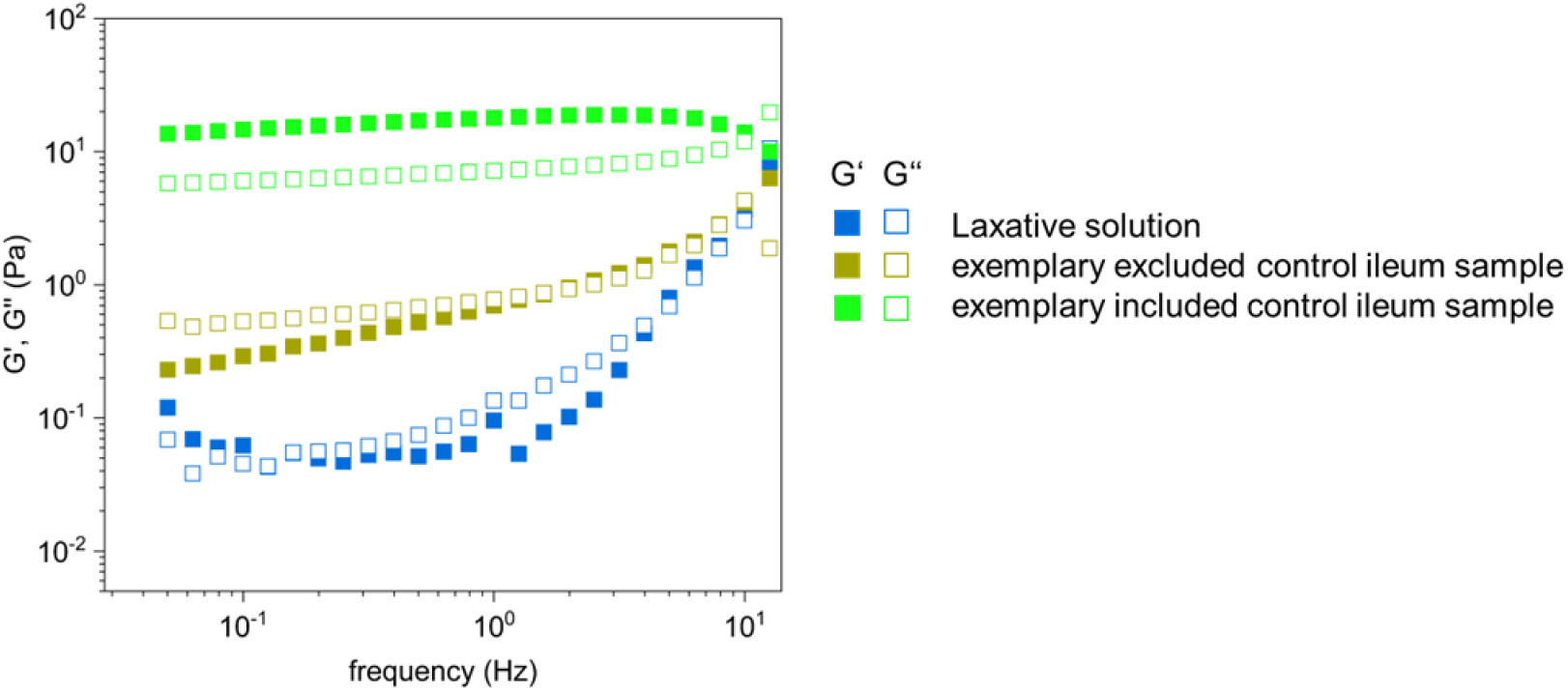
Comparison of storage modulus (G‘) and loss modulus (G‘‘) of an exemplary laxative solution (blue rectangles) and two ileal mucus samples from controls. The olive rectangles represent a sample of ileal mucus from the control group that appeared macroscopically diluted with the laxative solution and therefore had reduced viscoelastic properties; it was excluded from the analysis. To demonstrate the difference, we present a sample (in green rectangles) that was included in the analysis.

#### Ratio of G‘ to G‘‘ of ileal mucus from controls and Crohn’s Disease patients

**Supplementary Figure 2:**
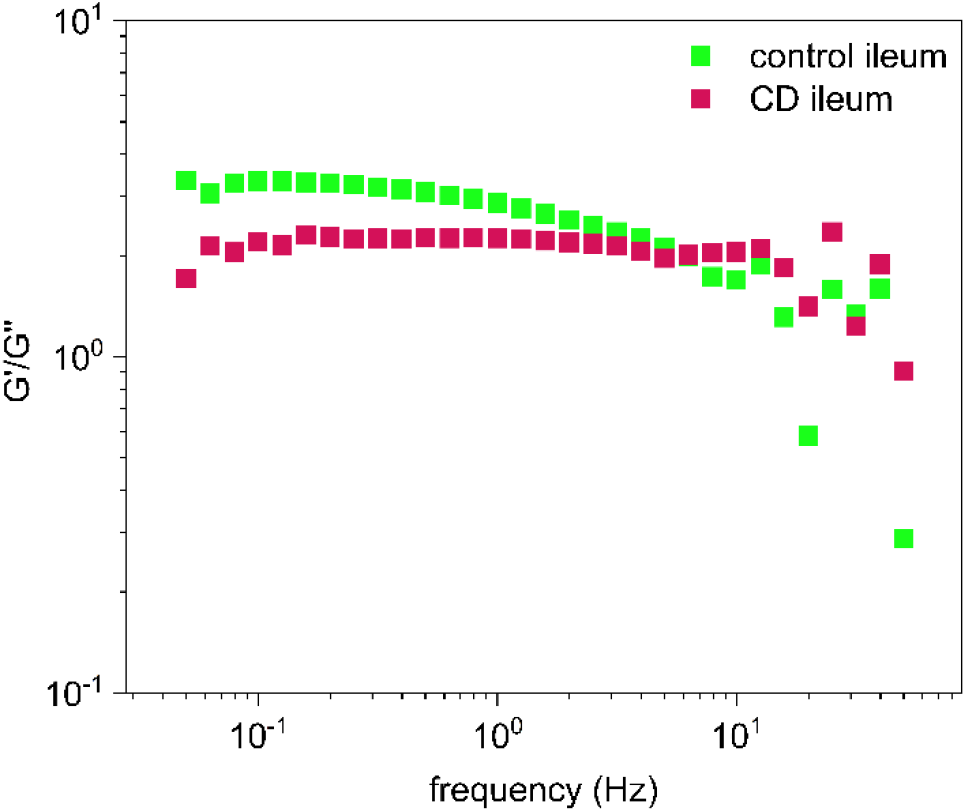
Quotient of the storage modulus G’ and the loss modulus G’’ of the mean value of ileal mucus from controls (n=11) and from Crohn’s disease (CD) patients (n=11), measured as a function of frequency. The quotient describes the relative elasticity of the samples, and its relatively constant course over the investigated frequency range indicates stable measurements.

#### No difference between samples that were measured on the same day or one day after collection

**Supplementary Figure 3:**
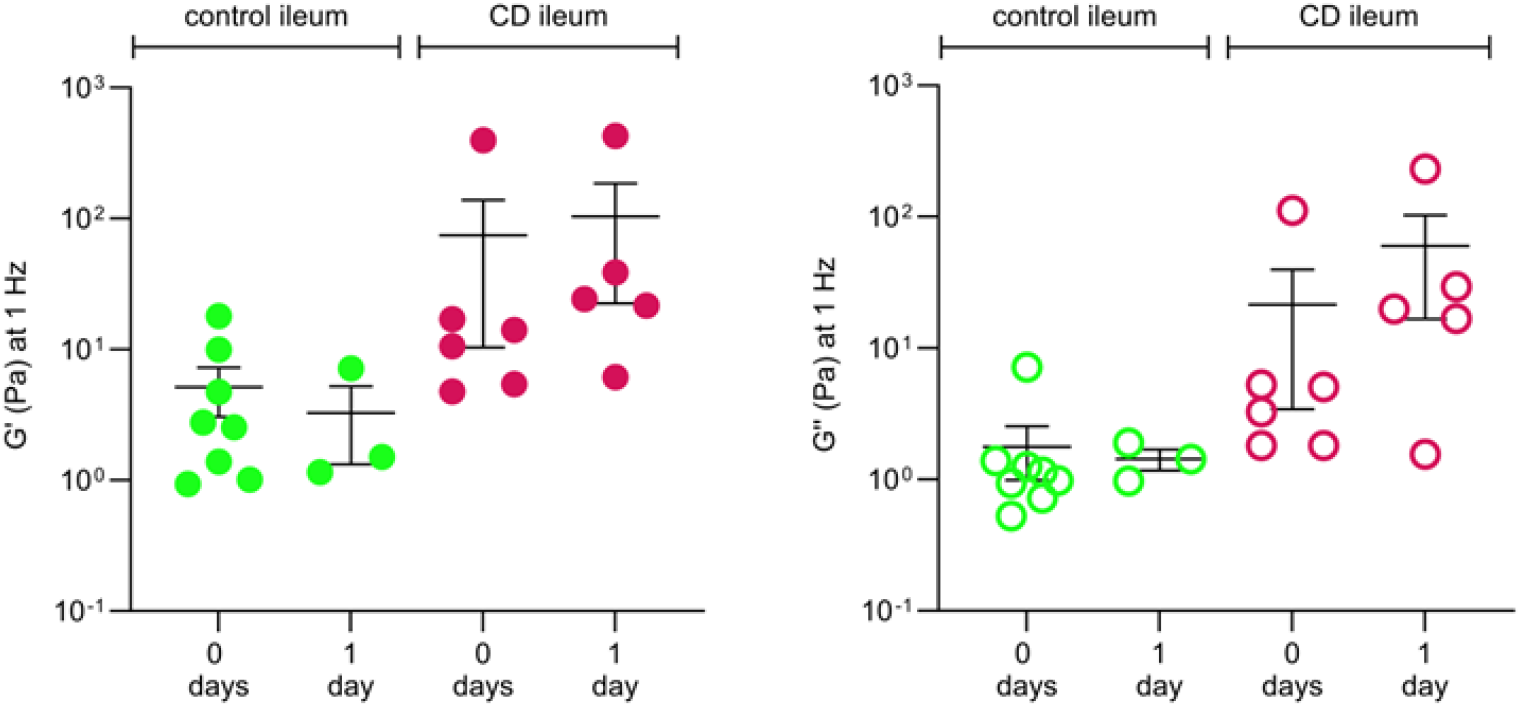
Storage modulus G’ and loss modulus G’’ of ileal mucus from controls (n=11) and from Crohn’s disease (CD) patients (n=11) were measured at a frequency of 1 Hz, either on the day of collection (0 days) or one day after (1 day). Data are shown as individual values and with the mean +/− standard error of the mean (SEM). These data indicate that different storage times are not the reason for the differences observed between mucus from controls and CD patients.

#### Airway mucus from controls that was used as reference showed constant frequency sweeps

**Supplementary Figure 4:**
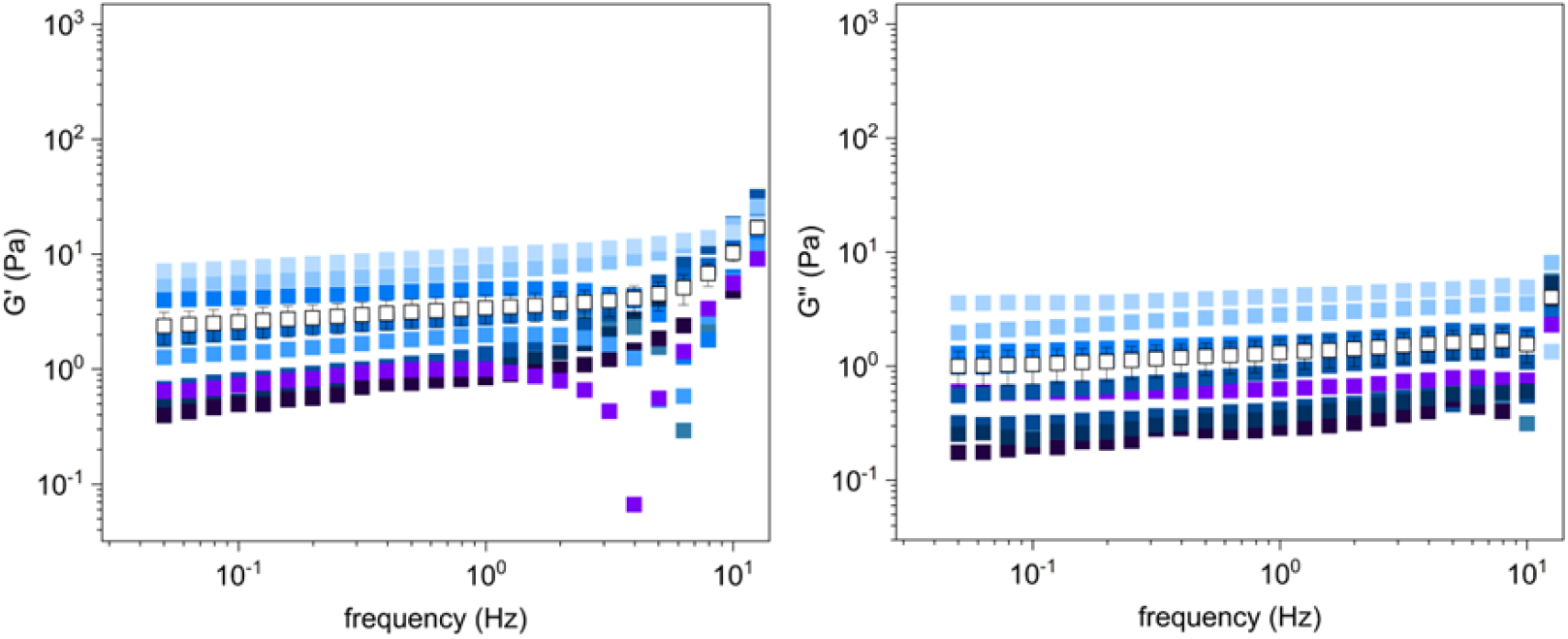
Storage modulus G’ and loss modulus G’’ of airway mucus from controls (n=10) were measured as function of frequency. Data are shown as individual values and with the mean +/− standard error of the mean (SEM) (open black squares with error bars).

#### Rheological data related to different clinical characteristics of controls and Crohn’s Disease patients

**Supplementary Table 2:**
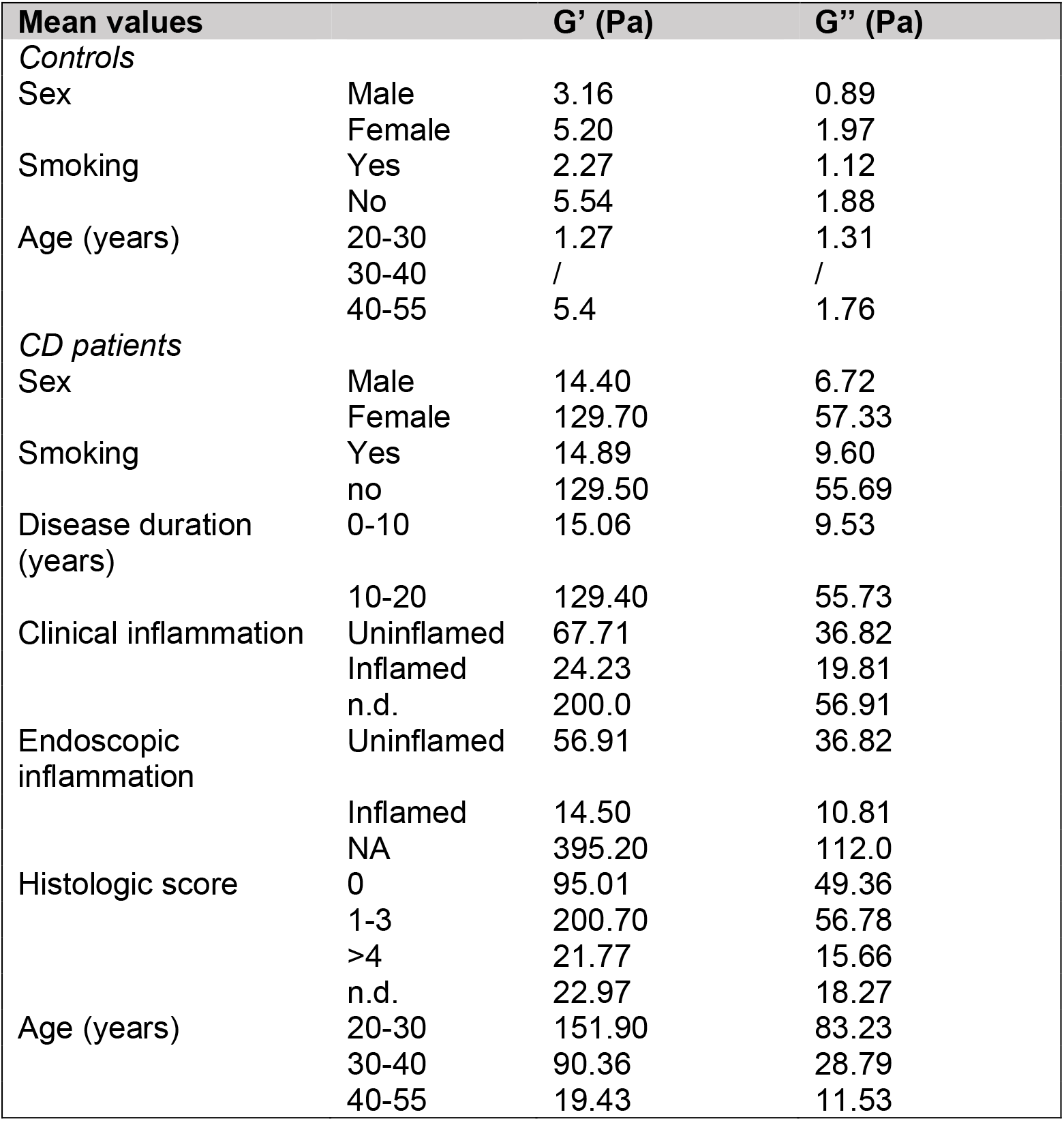
Storage modulus G’ and loss modulus G’’ of ileal mucus from controls (n=11) and Crohn’s disease (CD) patients (n=11) measured at a frequency of 1 Hz. The data are shown as the mean value categorized by different clinical characteristics shown in Figure 3. Abbreviation: n.d. = not determined.

#### The rheological behavior of mucus from controls and Crohn’s disease patients do not show any age dependency

**Supplementary Figure 5:**
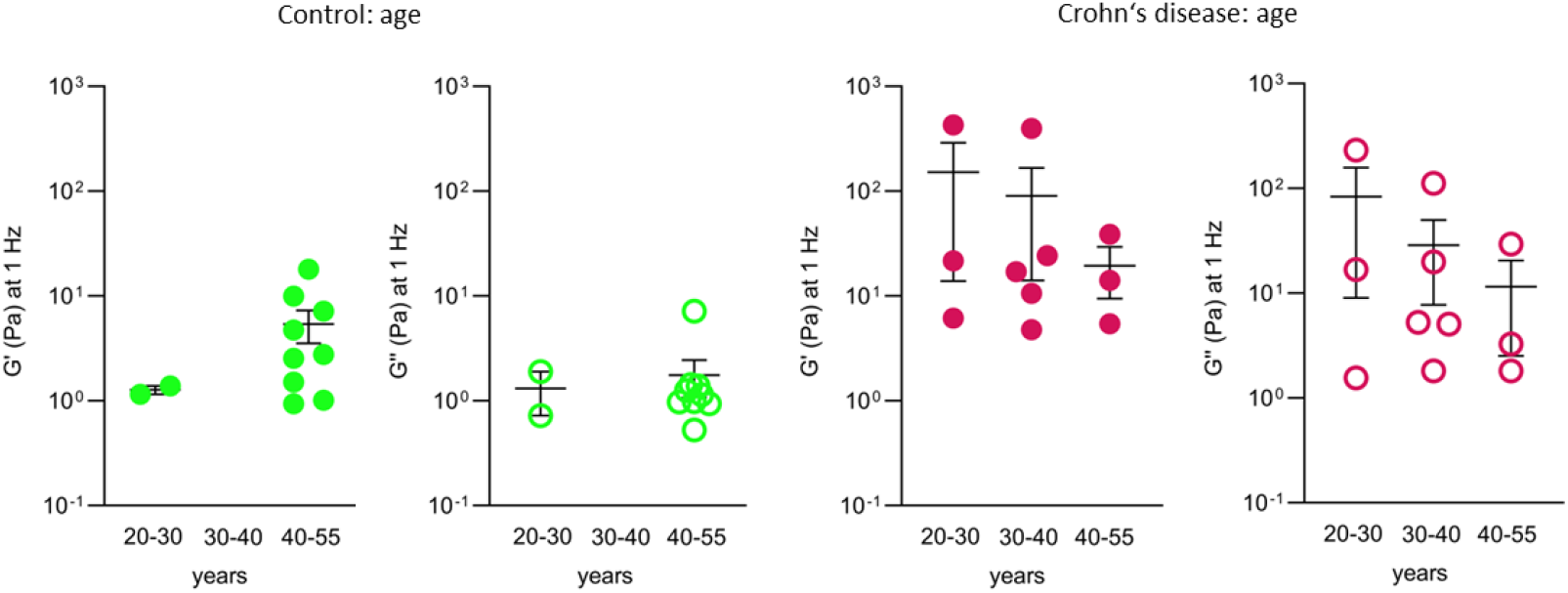
Storage modulus G’ and loss modulus G’’ of ileal mucus from controls (n=11; green squares) and Crohn’s disease patients (n=1; red squares1) were measured at a frequency of 1 Hz and divided depending on age (years). Data are shown as individual values and with the mean +/− standard error of the mean (SEM) and show no significant difference between age groups.

